# Evolutionary dynamics of asexual hypermutators adapting to a novel environment

**DOI:** 10.1101/2021.07.28.454222

**Authors:** Wei-Chin Ho, Megan G. Behringer, Samuel F. Miller, Jadon Gonzales, Amber Nguyen, Meriem Allahwerdy, Gwyneth F. Boyer, Michael Lynch

## Abstract

How microbes adapt to a novel environment is a central question in evolutionary biology. While adaptive evolution must be fueled by beneficial mutations, whether higher mutation rates facilitate the rate of adaptive evolution remains unclear. To address this question, we cultured *Escherichia coli* hypermutating populations, in which a defective methyl-directed mismatch repair pathway causes a 140-fold increase in single-nucleotide mutation rates. In parallel with wild-type *E. coli*, populations were cultured in tubes containing Luria-Bertani broth, a complex medium known to promote the evolution of subpopulation structure. After 900 days of evolution, in three transfer schemes with different population-size bottlenecks, hypermutators always exhibited similar levels of improved fitness as controls. Fluctuation tests revealed that the mutation rates of hypermutator lines converged evolutionarily on those of wild-type populations, which may have contributed to the absence of fitness differences. Further genome-sequence analysis revealed that, although hypermutator populations have higher rates of genomic evolution, this largely reflects the effects of genetic draft under strong linkage. Despite these linkage effects, the evolved populations exhibit parallelism in fixed mutations, including those potentially related to biofilm formation, transcription regulation, and mutation-rate evolution. Together, these results generally negate the presumed relationship between high mutation rates and high adaptive speed of evolution, providing insight into how clonal adaptation occurs in novel environments.

**Significance statement:** While mutations are critical source for the adaptation in a new environment, whether or not the elevated mutation rates can empirically lead to the elevated adaptation rates remains unclear, especially when the environment is more heterogenous. To answer this question, we evolved *E. coli* populations with different starting mutation rates in a complex medium for 900 days and then examined their fitness and genome profiles. In the populations that have a higher starting mutation rate, despite faster genome evolution, their fitness improvement is not significantly faster. Our results reveal that the effect of elevated mutation rates is only very limited, and the mutations accumulated in hypermutators are largely due to linkage effect.

## Introduction

Beneficial mutations are the ultimate source of adaptive evolution. Therefore, it is of interest to study how changes to mutational processes can influence an adaptive process. In terms of mutation rates, a theoretically complicated relationship with the rate of adaptation in asexual populations was proposed in the early studies on the evolution of sex (Muller 1932; Crow and Kimura 1965). These studies posit that, when mutation rates are low, such that the waiting time for a beneficial mutation to arise in a population remains long, increases in the mutation rate can result in linear increases in the adaptation rate. In contrast, when mutation rates are relatively high, such that multiple beneficial mutations frequently arise in different individuals within a population, beneficial mutations may interfere with each other’s opportunity to spread through the entire population. Thus, the facilitation effect of the mutation rate on the adaptation rate becomes diminished, a phenomenon later called clonal interference (Gerrish and Lenski 1998). More recent theoretical studies have suggested that the effective number of beneficial mutations per population is critical for the strength of clonal interference, and effective population sizes and effect-size distributions of mutations have also been proposed to be influential (Gerrish and Lenski 1998; Wilke 2004; Kim and Orr 2005; Bollback and Huelsenbeck 2007; Desai and Fisher 2007; Park and Krug 2007; Campos and Wahl 2010; Park et al. 2010; Good et al. 2012; Penisson et al. 2017). The relationship between mutation rates and rates of adaptation is further complicated because mutation rates can be plastically different in various environments (Williams and Foster 2012; Long et al. 2016; Shewaramani et al. 2017) and evolve over time (Wielgoss et al. 2013; Swings et al. 2017). Given that most mutations are deleterious, high mutation rates create high mutational loads in the genome, potentially driving the spread of antimutator alleles and resulting in the evolution of lower mutation rates (Muller 1950; Kimura 1967; Lynch 2008). According to the drift-barrier hypothesis, the reduction of the mutation rate should continue until the addition of new antimutator alleles no longer contributes to a significant enough reduction in mutational load to overcome genetic drift (Lynch 2010b; Sung et al. 2012; Lynch et al. 2016). Consequently, if two populations with initially different mutation rates adapt to the same constant environment, the resulting difference in fitness-improvement rates can be less than predicted if both populations converge evolutionarily to similar mutation rates.

While ample theoretical discussion exists on the relationship between mutation rates and the rate of adaptation, the theory has been mostly focused on constant environments with a simple fitness landscape, neglecting the likely complexity of more natural environments. The limited empirical evidence in asexual populations suggests that adaptation rates are a concave-down function of the mutation rate (Arjan et al. 1999; Desai et al. 2007; Sprouffske et al. 2018). However, the experimental environments in these studies were generally simple and homogeneous (Arjan et al. 1999; Desai et al. 2007; Sprouffske et al. 2018), or the populations of interest were already well-adapted to the experimental environment (McDonald et al. 2012). How the relationship between mutation rates and adaptation rates evolves when the environmental setting becomes more complex and heterogeneous is less understood. For example, the modes of adaptation in complex environments may vary greatly due to the presence of a more rugged fitness landscape, additional paths available for fitness improvement, the presence of multiple spatial or nutritional niches, or more complicated genetic interactions. (Handel and Rozen 2009; Lynch 2010a; Ochs and Desai 2015; Guo et al. 2019). Therefore, it is necessary to study adaptive processes in more complex and heterogeneous environments to determine whether the principles observed in simpler environments still apply.

To study how the mutation rate affects adaptation in a more complex setting, we performed long-term experimental evolution of *Escherichia coli* in culture tubes containing a complex medium, Luria-Bertani (LB) broth, which is comprised of a nutritionally-rich mixture of multiple amino-acid based carbon sources (Sezonov et al. 2007). In contrast to evolution in flasks containing glucose-limited media, such environments can facilitate the rapid emergence of stable subpopulations and clonal divergence based on spatial niche differentiation and amino-acid metabolism divergence (Behringer et al. 2018). To vary the mutation rate, we evolved both WT populations (MMR+) and hypermutator populations with an impaired methyl-directed mismatch repair pathway (MMR-, obtained by *mutL* knockout), for which the single-nucleotide mutation rate is 140-fold higher than that for the WT genetic background (Lee et al. 2012). Because the results of experimental evolution may be altered by different demographic settings (Vogwill et al. 2016; Wein and Dagan 2019), replicated populations were assigned to one of three different daily-transfer size treatments: 1/10 (large, L), 1/10^4^ (medium, M), and 1/10^7^ (small, S) dilutions into fresh media. Here, we examined the differences in phenotypic and molecular evolution among these populations over the course of 900 days.

## Results

### Higher initial mutation rates do not translate into faster rates of fitness improvement

When batch cultured, *E. coli* commonly adapt to their experimental environments and show fitness improvement compared to their ancestors (Van den Bergh et al. 2018; McDonald 2019). To compare adaptation rates in populations originating from genetic backgrounds with different initial mutation rates (MMR- and WT), we performed head-to-head competition assays between populations that had evolved for 900 days and their corresponding ancestor. For each of six genetic-background/transfer-size combinations (2 × 3), four replicated populations were measured. Across all 21 populations with data available (three were aborted; see Materials and Methods), mean fitness significantly increased relative to the time-zero ancestor, by a ratio of 1.14 (SE = 0.019; *P* = 5.8 x 10^-7^, two-tailed *t*-test), indicating the evolution of these populations shaped by adaptive processes.

The amount of fitness improvement of MMR-populations was not significantly different from that for WT populations among any of the transfer sizes (**Fig. 1A**; L: *P* = 0.26; M: *P* = 0.46; S: *P* = 0.15, nested ANOVA). In particular, considering the ratio of mean fitness improvement (MMR- : WT), no transfer size produced a ratio significantly different from 1.0. For example, for the L transfer size, the mean fitness improvement for MMR- and WT backgrounds are respectively 0.27 (SE = 0.043) and 0.21 (SE = 0.047), and therefore the ratio (MMR-: WT) is 1.28 (SE = 0.35). Similarly, in the M and S transfer sizes, the ratios are 0.69 (SE = 0.28) and 2.76 (SE = 1.98), respectively. Thus, starting evolution as a hypermutator does not necessarily translate into a faster fitness-improvement rate.

**Figure 1.**
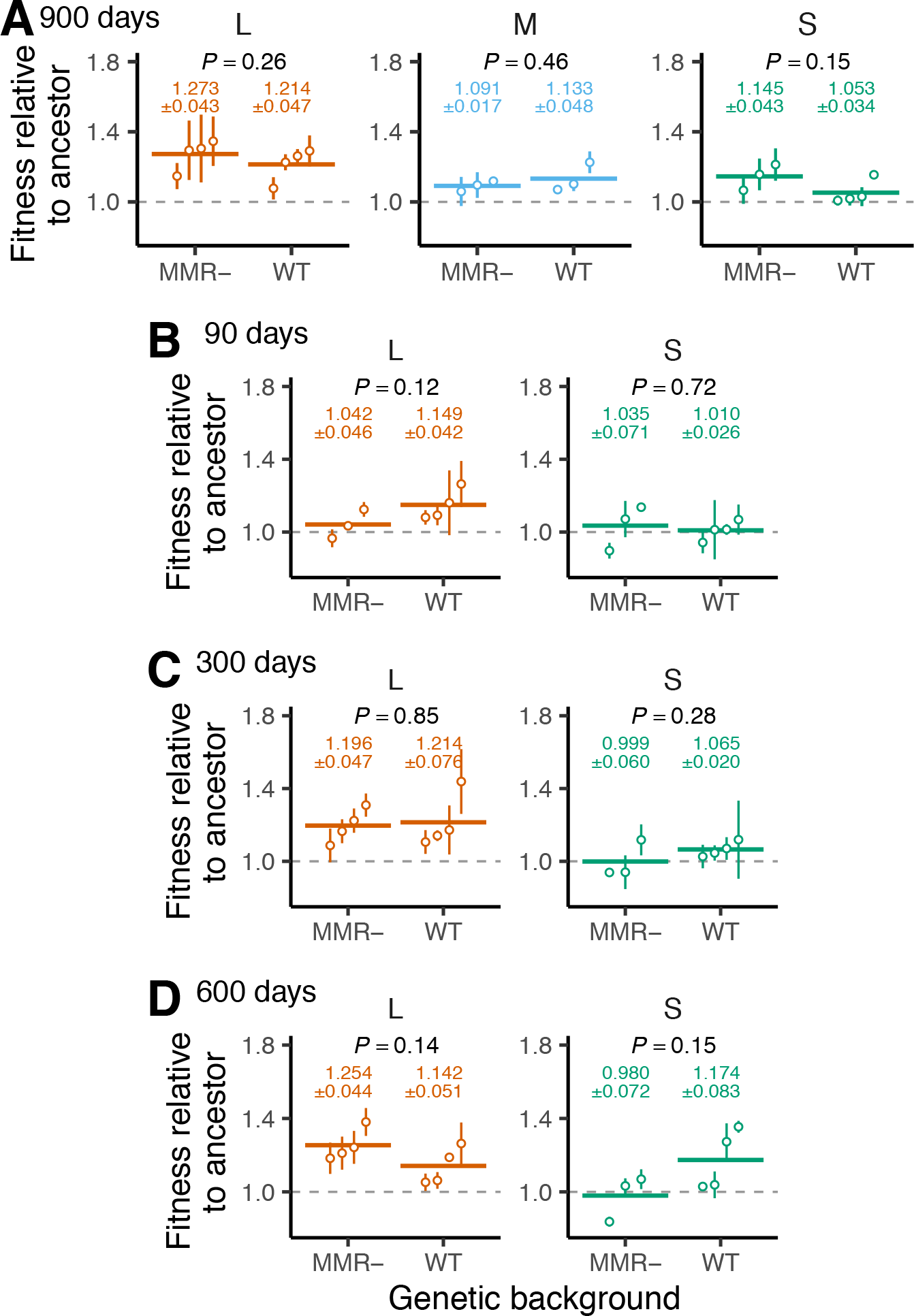
Fitness improvement during experimental evolution. For evolved populations under different transfer sizes (orange for L, blue for M, or green for S) and different genetic backgrounds (MMR- or WT), mean fitnesses relative to the ancestor at **(A)** day 900, (**B**) day 90, **(C)** day 300, or **(D)** day 600 are reported. Each open circle represents an estimated mean for an evolved population with at least three independent competition assays. The error bars represent SEs. Gray dashed lines represent no improvement from ancestral fitness. The means across all evolved lines for a combination of transfer size and genetic background are represented by colored horizontal lines; the numeric values of means and SEs are printed on the top. The *P*-values for nested ANOVA are also shown on the top.

Previous studies of *E. coli* in simpler, more homogeneous environments have shown that the most rapid increases in population fitness typically occur within 2500 generations, after which the rate of adaptation significantly slows (Barrick et al. 2009). At 900 days, the L populations, which due to their large transfer size experienced the least number of cell divisions, had experienced ∼3000 generations, whereas the S populations had experienced ∼21,000 generations. As such, the absence of significant differences in the cumulative amount of adaptation over this period – despite large differences in initial mutation rate – might be a consequence of both genetic backgrounds having exited an initial period of rapid fitness evolution. Thus, to better survey any temporal heterogeneity in the rate of adaptation, we further assessed fitness after 90-, 300-, and 600-days of evolution in response to L and S transfer sizes. The results of nested ANOVA again demonstrate a lack of evidence for an increase in initial mutation rates leading to an increase in the amount of fitness improvement (**Fig. 1B-D**). The results are not qualitatively changed even when the natural-logarithmic transformed fitness is used in the analysis (**Fig. S1**). Thus, high mutation rates did not result in accelerated fitness improvement in these asexual populations even in the early stages of adaptation.

### Mutation rates evolve to be more similar throughout experimental evolution

The indifference of adaptation rates to initial mutation rates might be explained if the mutation rates of hypermutator and WT populations became more similar during the evolution experiment, either due to a reduction in the mutation rate in initially hypermutating populations or to an increase of the rate in WT populations. To test this possibility, we performed fluctuation tests, which indirectly measure mutation rates at a resistance locus (Foster 2006), on different clones isolated from evolved populations after 900 days. Although the ratio of rifampicin-resistance mutation rates for the two ancestral lines (MMR- : WT) was 250 (**Fig. S2**), after 900 days of evolution, the mean difference in mutation rates between MMR- and WT backgrounds greatly decreased across all transfer sizes (**Fig. 2**). For example, in the L transfer size, the mean mutation rate is 5.0 x 10^-7^ (SE = 2.7 x 10^-7^) and 1.6 x 10^-8^ (SE = 1.2 x 10^-8^) for MMR- and WT evolved populations, respectively. Therefore, the ratio of mean mutation rates (MMR- : WT) was reduced to 32 (SE = 29). This reduction, however, seems mostly attributed to the occasional emergence of higher mutation rates in the WT background as only one population (115) among four tested WT, L populations shows a significant increase of the mutation rate in both tested clones. In the M and S transfer size, the ratio of mean mutation rates (MMR- : WT) was also reduced to 32 (SE = 15) and 12 (SE = 3.9), respectively. However, the repeated evolution of lower mutation rates in clones isolated from MMR-background plays a more important role in the reduction as all of the tested clones from MMR-, M or S populations show a significant decrease of the mutation rate. Thus, the evolutionary convergence of mutation rates likely contributes to the difference in adaptation rates being less than what might be expected based on initial differences in mutation rates, but interestingly, the specific evolutionary mechanisms underlying these similarities are different. These observations demonstrate how transfer schemes can affect the evolutionary dynamics of mutation rates in asexual populations.

**Figure 2.**
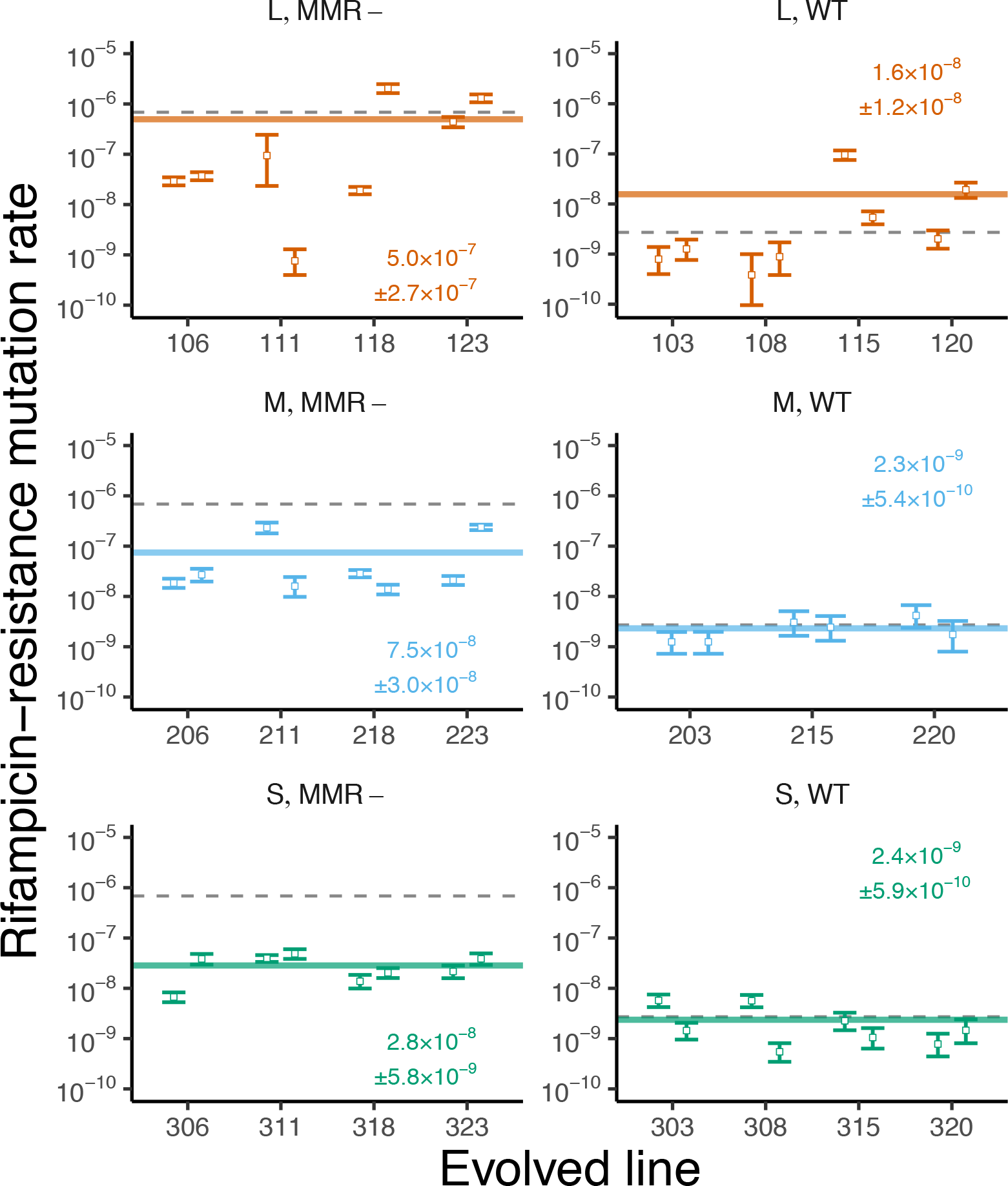
Evolution of mutation rates after 900 days of experimental evolution. Each panel shows mutation rates of evolved populations in different combinations of transfer sizes (L, M, or S) and genetic background (MMR- for mismatch repair defective or WT for wild-type). In each combination, three or four evolved populations were tested. Two clones per evolved population were isolated and measured. The open circles and error bars represent the mean and the 95% confidence interval for each clone. The grey dashed line represents the mutation-rate measurement of the corresponding ancestor. The colored horizontal lines represent the mean mutation-rate measurement of each combination; the value of means and their SEs are also printed.

### Genome evolution rates are less different than predicted by initial mutation rates

To enhance our understanding of how the tempo and mode of genomic evolution relate to fitness and phenotypic evolution, we performed metapopulation sequencing of each experimental population roughly every 100 days to acquire mutation profiles of associated derived allele frequencies (DAFs). For each combination of genetic background and transfer size, we estimated the rate of genomic evolution by regressing the number of mutations per clone (i.e., the sum of DAFs of all observed SNPs) of all populations against the number of generations at each sampled time point. Because a quadratic regression model did not significantly outperform the linear regression (*P* = 0.028 for MMR-/L; *P* > 0.10 for the others, nested ANOVA), we will focus on the results of the linear regression below.

As with the rate of adaptation, for all three transfer sizes, the ratio of the rate of genomic evolution between the two backgrounds (MMR-: WT) was much smaller than the initial difference in mutation rates (**Fig. 3A**). For example, under the L transfer size, the rate of genomic evolution is 115 (SE = 5.3) and 30 (SE = 4.3) mutations per clone per 1000 generations for MMR- and WT populations, respectively. Therefore, the ratio of the genomic-evolution rates is only 3.8 (SE = 0.58). Similarly, in the M and S transfer sizes, the ratio is 20 (SE = 1.3) and 23 (SE = 1.6), respectively. This observation remained qualitatively similar even when different kinds of measurements for genomic divergence were used, e.g., the number of detected mutations (**Fig. S3A**), nucleotide diversity (**Fig. S3B**), or when genomic divergence was estimated only by synonymous SNPs (**Fig. S4**).

**Figure 3.**
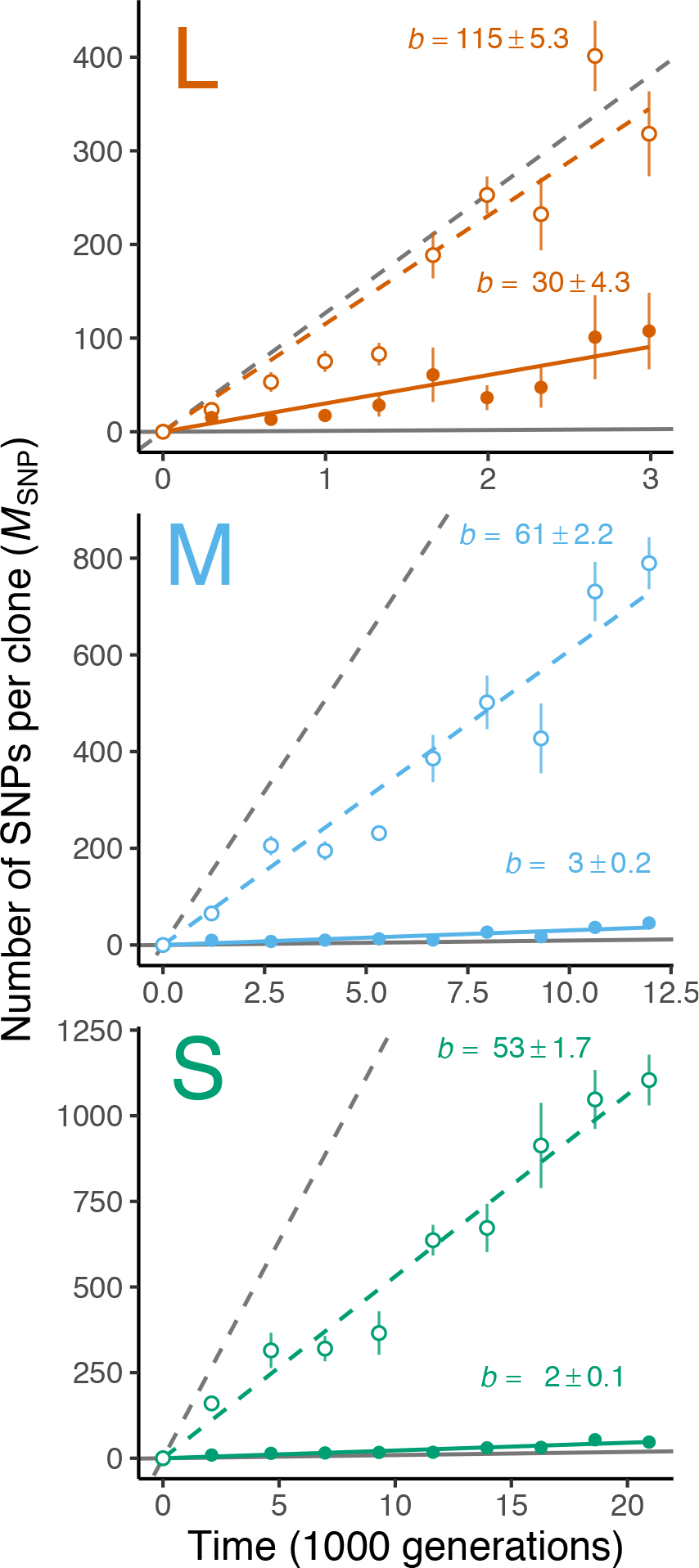
Rate of genomic evolution in evolved populations. Each panel shows the results of evolved populations in two genetic backgrounds (MMR- and WT) in a transfer size (L, M, or S). Each dot shows a mean number of SNPs per clone for MMR- (open circles) or WT (closed circles) populations at a sequencing time point. The error bars represent the associated standard errors. The dashed and solid lines are linear regressions against the time for MMR- and WT populations, respectively. The estimated slope (*b*) and associated standard error are also printed for each regression line. The grey dashed and solid lines represent how evolved populations are expected to accumulate mutations based on the initial mutation rates of MMR- and WT ancestors.

To further survey how genomic evolution rates vary across experimental populations and to determine if the mean rates accurately represent the majority of experimental populations, we separately measured the rate of genomic evolution for each experimental population (**Fig. S5**). This revealed that the distribution of rates in the WT populations under the L transfer size is wider than the distributions of rates in all other combinations of genetic background and transfer sizes. Specifically, in the L transfer size, although five of the eight WT populations exhibit a genomic evolution rate of ∼10 mutations per clone per 1000 generations, one WT population (115) has a rate about 2× higher, and two WT populations (101 and 113) have a rate close to 100, similar to MMR-populations. Consistent with the results of fluctuation tests noted above, these results suggest that some, but not all, WT populations under the L transfer size evolved a higher mutation rate (see Discussion).

### Mutations arising in hypermutators are more likely to be fixed

Using longitudinal metagenomics-sequencing data allows one to observe evolutionary dynamics at the level of single mutations and thus better understand the entire adaptive process. Here, we will focus on fixed mutations because they are more likely to contribute to adaptive processes than polymorphic mutations or other mutations that are transient in a population. Because our experimental-environment setting facilitates the development of subpopulation structure, we applied clade-aware hidden Markov chain (caHMM) analysis. Assuming coexistence of two clades (major and minor) in the population, caHMM considers each mutation’s DAFs found in different sequencing time-points and then infers which clade each mutation belongs to, whether each mutation reaches within-clade fixation, and when a fixed mutation reaches fixation (Good et al. 2017).

One characteristic parameter in an adaptive process is the probability that mutations reach fixation in a population. With the observed numbers of both detected and fixed mutations, we first tested whether hypermutator populations have a different probability of mutation fixation within clades compared to WT populations. Different mutation types have a different potential to impact populational fitness. For example, compared to synonymous SNPs, nonsynonymous SNPs have a greater potential to change protein functions, intergenic SNPs have a greater potential to change protein expression, and structural variations (SVs; including indels and mobile-element insertions) have a greater potential to disrupt a protein. Therefore, we performed separate tests on the conditional fixation probability for these four functional categories of mutations.

While not always significant, the fixation probability in the MMR-populations is generally higher than in WT populations across different transfer sizes and different categories of mutations (**Fig. 4A**). The fixation probability is similar across different categories of mutations, regardless of their perceived potential to affect fitness, suggesting that the fixation of most mutations is a consequence of genetic hitchhiking as opposed to intrinsic beneficial effects.

**Figure 4.**
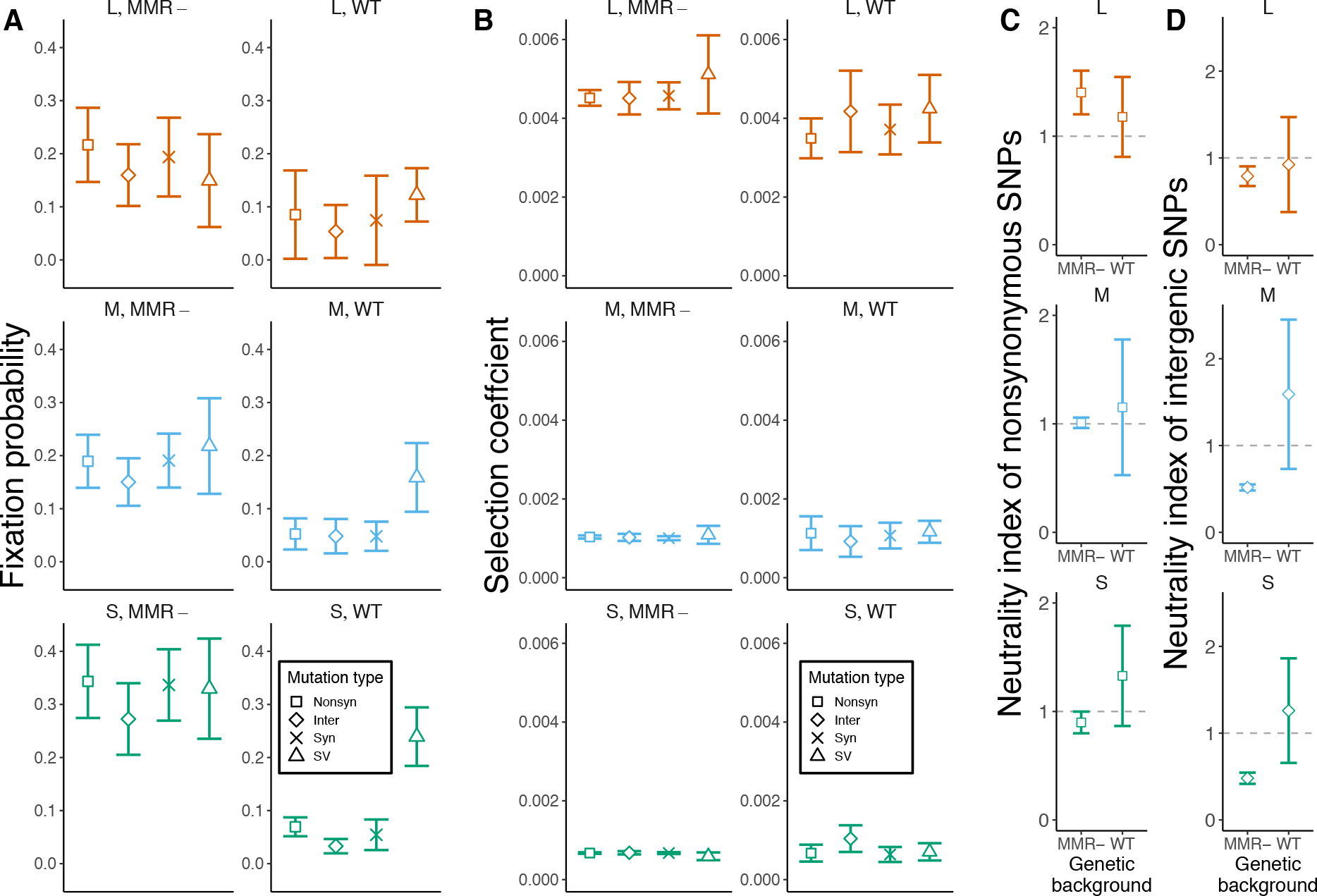
Analysis of strength of natural selection associated with fixed mutations in different categories in different treatments of experimental evolution. **(A)** Each symbol shows the population mean probability of <u>nonsyn</u>onymous mutations (squares), <u>inter</u>genic mutations (diamonds), <u>syn</u>onymous mutations (crosses), or structure variation mutations (SV; triangles) that reached within-clade fixation in each combination of transfer size (L, M, or S) and genetic background (MMR- or WT). The error bars show the 95% confidence intervals. **(B)** Each symbol shows the mean selection coefficient of <u>nonsyn</u>onymous mutations (squares), <u>inter</u>genic mutations (diamonds), <u>syn</u>onymous mutations (crosses), or SV (triangles) that are fixed in any clade in any population belonging in a combination of transfer size and genetic background. The error bars show the 95% confidence intervals. **(C)** Each square shows the population mean neutrality index of nonsynonymous mutations for a combination of transfer size and genetic background. The error bars show the 95% confidence intervals. The grey lines indicate where the value = 1.0. (**D**) Each square shows the population mean neutrality index of intergenic mutations for a combination of transfer size and genetic background. The error bars show the 95% confidence intervals. The horizontal grey lines denote the point of neutrality (1.0).

Another critical factor determining the temporal dynamics of a mutation is the underlying fitness effect. With the temporal data of allele frequencies for a mutation, we can quantify the net selection coefficient, which reflects its own fitness effects but can be potentially affected by the effects of linked mutations. We did not find a significant difference between the selection coefficients of fixed mutations in the two genetic backgrounds (**Fig. 4B**). More importantly, we also found that different categories of mutations show similar mean selection coefficient estimates, which again suggests a pivotal contribution of hitchhiking effects to the mutational dynamics of genome evolution these in asexual populations.

### Neutrality tests reveal partial evidence for positive selection

In theory, comparing the number of fixed mutations in functional categories of sites with different potential effects on fitness can summarize general patterns in the mode of genome evolution (McDonald and Kreitman 1991; Rand and Kann 1996). If a population has experienced strong positive selection on protein function or expression, it is expected that there will be more fixed mutations with a greater potential to change protein function or expression than mutations with smaller such potential. If a population has experienced purifying selection on protein function or expression, it is expected that there will be less fixed mutations with a greater potential to change protein function or expression than mutations with smaller such potential. Therefore, the ratio of the number of fixed nonsynonymous synonymous SNPs (*F*_N_) to the number of fixed synonymous SNPs (*F*_S_) or the ratio of the number of fixed intergenic SNPs (*F*_I_) to *F*_S_ is predicted to be large with a strong positive selection. Similarly, *F*_N_/*F*_S_ or *F*_I_/*F*_S_ are predicted to be small with a strong purifying selection. However, these ratios are not directly comparable in MMR- and WT backgrounds, because the two genetic backgrounds have varied mutational spectra and different relative rates of nonsynonymous and synonymous mutations (Lee et al. 2012). To address this issue, we normalized the observed *F*_N_/*F*_S_ by the ratio of nonsynonymous and synonymous mutations (*U*_N_/*U*_S_) previously observed in a mutation accumulation experiment which utilized our exact ancestral genotypes (Lee et al. 2012). The ratio of these ratios, (*F*_N_/*F*_S_)/(*U*_N_/*U*_S_), will be referred to as the neutrality index of nonsynonymous SNPs, since being significantly > 1.0 or < 1.0 implies a predominance of positive selection in driving mutation fixation or a predominance of purifying selection in driving mutation extinction. It is analogous to Tachida’s index for neutrality (Tachida 2000) but with a different normalizing approach – *via* results from mutation-accumulation experiment instead of polymorphism. This definition of the neutrality index can be applied to any kind of mutations. For example, we also define the neutrality index of intergenic SNPs as (*F*_I_/*F*_S_)/(*U*_I_/*U*_S_), where *U*_I_ is the number of intergenic mutations observed in mutation accumulation (Lee et al. 2012).

The neutrality index of nonsynonymous SNPs in MMR-populations under the L transfer size is significantly larger than one, consistent with the model of strong positive selection (**Fig. 4C**). Moreover, the neutrality index of intergenic SNPs in MMR-populations under all transfer sizes is significantly smaller than one, suggesting overall strong purifying selection on intergenic SNPs (**Fig. 4D**). On the contrary, the neutrality index in WT populations is never significantly different from one. While this can indicate a weaker strength of selection in WT populations, it may also be the result of insufficient statistical power due to an overall smaller number of fixed SNPs in WT populations, evident by their large confidence intervals associated with the point estimation.

### Parallel evolution of fixed mutations at the genic and nucleotide level

As the previous analysis suggests that the statistical power of the neutrality index to reveal positive selection may be insufficient, we also considered parallelism of fixed mutations to examine the action of positive selection, using two different metrics. First, we utilized a sum of *G*-scores to measure the excess parallelism across fixed nonsynonymous mutations relative to the expectation based on gene lengths (Tenaillon et al. 2016). For each gene, a higher *G*-score implies that the gene is more enriched for fixed nonsynonymous mutations than expected by gene length. We then summed the *G*-scores of all genes for each combination of genetic background and transfer size and found statistical significance in all cases (*z*-test, **Fig. 5A**). Second, we calculated the mean Bray-Curtis similarity of the number of fixed nonsynonymous mutations across all genes (Turner et al. 2018) for all pairwise comparisons of evolved populations in each combination of genetic background and transfer size. The results again show statistical significance for similarity in all cases (*z*-test, **Fig. S6**). Therefore, both findings suggest that positive selection has shaped the genomic evolution of populations in all genetic-background / transfer-size combinations.

**Figure 5.**
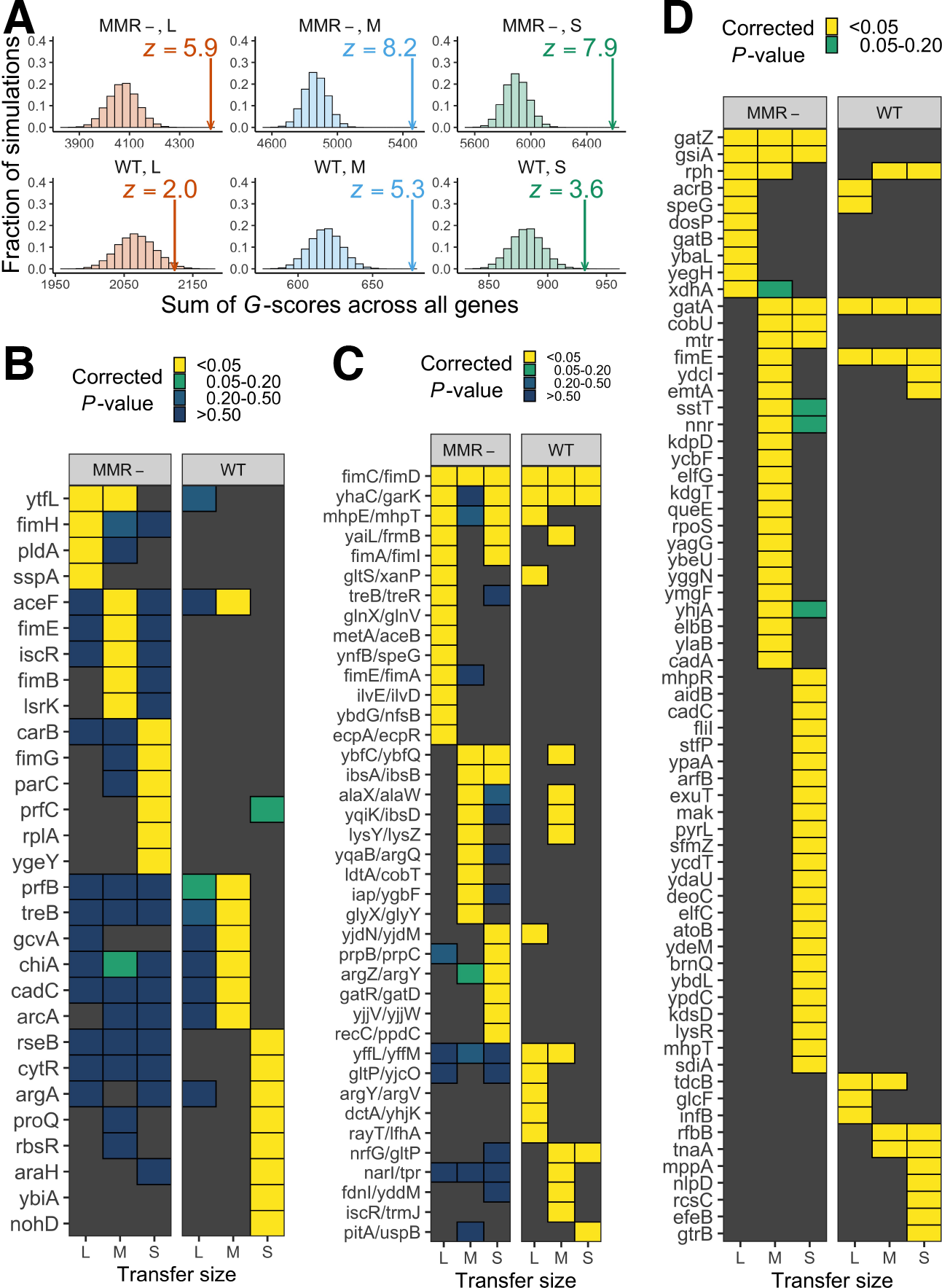
Evolutionary parallelism as evidence for positive selection. **(A)** The arrow and vertical dashed line show the observed sum of *G*-scores, representing the extent of parallel mutation for a particular combination of transfer size and genetic background. The histogram shows the distribution of 20,000 simulated sums of *G*-scores, representing the null distribution of evolutionary parallelism. The significance of the observed sums can be evaluated by *z*-scores (*z* > 1.65 for one-tailed *P* < 0.05). **(B)** List of genes with fixed nonsynonymous mutations. Significance levels (simulated *P*-values with Bonferroni correction) are shown by the different non-black colours of tiles. Genes with no such hits in a particular combination are shown by black tiles. **(C)** List of genes with fixed intergenic mutations. Significance levels (simulated *P*-values with Bonferroni correction) are shown by the different non-black colours of tiles. Intergenic regions with no such hits in a particular combination are shown by black tiles. **(D)** List of genes significantly overrepresented for structural mutations that are likely under positive selection. Yellow tiles highlight genes observed in at least two populations and yielding simulated *P*-values < 0.05 after Bonferroni correction.

The analysis of parallelism also helps reveal which mutations are most likely to be drivers of adaptation. Using the same *G*-score analysis, in each genetic-background/transfer-size combination, we identified genes that were overrepresented for fixed nonsynonymous mutations (*P* < 0.05, Bonferroni correction; **Fig. 5B**). Gene Ontology (GO) analysis on these gene subsets revealed that significantly enriched GO terms were often related to transcription regulation and biofilm formation (**Table S1**). Using similar methods, we also identified subsets of genes enriched for fixed mutations in intergenic regions (**Fig. 5C**) and for structural mutations, including indels and IS-element insertions (**Fig. 5D**). Together, the genes in these lists serve as good candidates for revealing the various mechanisms important to adaptation in complex environments (see Discussion).

In addition to genic-level parallelism, we also identified nucleotide-level parallelism. In particular, 197 cases of parallel fixed nonsynonymous mutations were identified in at least two experimental populations within the same combination of transfer size and genetic background (**Table S2**). Because the probability that fixed mutations would occur by chance at a given nonsynonymous site in at least two experimental populations is very low, the observed parallelism at the nucleotide-level again suggests positive selection and identifies important candidates for studying the molecular mechanisms associated with adaptation in complex media. For example, three instances of parallel mutation within *fimH* were located in its mannose-binding domain, and therefore may be good candidates for future functional studies of cell-adhesion (Schembri et al. 2001). Furthermore, three instances of parallel-fixed nonsynonymous mutations were found in genes with GO terms associated DNA repair or DNA replication, including *dnaE* (DNA pol III subunit α), *yajL* (protein/nucleic acid deglycase 3), and *nrdA* (ribonucleoside-diphosphate reductase 1, α subunit dimer). As all three of these cases arose in M and S transfer sizes with MMR-backgrounds, they may be good candidates for studying the molecular mechanisms associated with lowering mutation rates.

## Discussion

Here, we describe the effect of the initial mutation rate on the rate of fitness improvement and genomic evolution by experimentally evolving *E. coli* with two distinct mutation-rate backgrounds. While the initial difference in mutation rates of these two genetic backgrounds is >100×, after 900 days of evolution, the differences in the net rates of fitness improvement (0.7-2.7 fold), in the mutation rates (12-32 fold), and in level of genome evolution (4-23 fold) are much smaller. At the resolution of single mutations, we found that mutations arising in MMR-populations exhibit higher fixation probabilities than mutations in WT populations, while mutations in different categories with presumably different fitness effects show similar fixation probabilities within each genetic-background treatment, suggesting the strong influence of hitchhiking effects in genome evolution in these populations. Despite the strong linkage and pervasive genetic draft, there is still evidence, in the form of a high degree of parallelism in mutations arising in particular genes and nucleotides, that positive selection shapes genome evolution in all transfer-size/genetic-background combinations. The observed mutations with high parallelism also serve as excellent candidates for understanding the mechanisms of adaption to the complex conditions provided by the experimental environment.

### Effects of high mutation rates on evolution

Our experiments reveal that, even for 900 days of evolution in a complex medium, hypermutating *E. coli* does not necessarily exhibit a faster rate of adaptation than wild-type *E. coli*. This observation is consistent with previous experimental-evolution results over a shorter period and in simpler media, such as glucose medium DM25 for 1000 generations (Arjan et al. 1999) and glucose medium DM1000 for 3000 generations (Sprouffske et al. 2018). Our experiments further demonstrate that populations with initially hypermutator backgrounds can rapidly evolve lower mutation rates. Together with other empirical work on prokaryotic (Sprouffske et al. 2018) and eukaryotic hypermutators (McDonald et al. 2012), these results suggest that the strong genetic load due to deleterious mutations remains a pivotal factor in the evolution of mutation rates, consistent with the drift-barrier hypothesis (Lynch 2010b; Sung et al. 2012). Even when some new hypermutator alleles can spread in a population by linkage with other beneficial mutations during the adaptive process and thus briefly improve the rate of adaptation (Sniegowski et al. 1997; Tenaillon et al. 1999; Tenaillon et al. 2001), such events are usually transitory (Giraud et al. 2001; Desai and Fisher 2007; Wielgoss et al. 2013). Moreover, the fact that lowering the mutation rate could not have involved a reversion of deleted MMR in our experiments implies that there is excess capacity for improving replication fidelity through other parts of *E. coli* genome. As mutation-rate evolution occurred over a relatively short period in our study, the results bear on several critical questions for future studies, including the rapidity of the dynamics of the evolution of mutation rates and the consequences for the mutational spectrum.

### Effect of genetic linkage in evolution

In asexual populations with reduced recombination, the fate of a mutation is largely affected by its association with other mutations due to strong genetic linkage (Gillespie 2000; Neher 2013; Couce et al. 2017). Extensive hitch-hiking is a feature of our evolving *E. coli* populations, as we observe similar fixation probabilities and net associated fitness effects for fixed mutations across different functional categories of mutations. To better illustrate this effect of genetic linkage, we showed that the temporal DAF changes of any two SNPs in the same genome are highly correlated (i.e., a rightly-skewed distribution of correlation coefficients) and result in more largely positive correlation coefficients than for a random expectation of non-linked mutations (**Fig. S7**). Accordingly, pre-existing non-beneficial mutations can become fixed by hitchhiking with newly-arising beneficial mutations, and these linkage effects can limit the rate of adaptation (Schiffels et al. 2011; Kosheleva and Desai 2013). Therefore, even though MMR-populations show a range of genome-evolution rates 4 -23× higher than WT populations, the excess of fixed mutations does not directly contribute to adaptation rates.

### Effects of transfer sizes in evolution

The three different transfer sizes (L, M, and S) implemented in our evolution experiment allow us to compare adaptive processes in different population-genetic environments. In theory, when the transfer size is large, reduced genetic drift renders a higher efficiency of promotion of beneficial mutations in a population. Consistent with this theoretical prediction, the populations cultured with the L-transfer size show highest rates of fitness gain (**Fig. 1**), for nonsynonymous mutations, the observed neutrality index of MMR-populations under L transfer size is lowest and < 1.0 (**Fig. 4C**). Compared to L-transfer treatment, the populations cultured with the S-transfer size may have accumulated more deleterious mutations and thus show less fitness improvement.

We also found that the populations under the L-transfer size tend to evolve higher mutation rates (**Fig. 2**), as hypermutator alleles may hitchhike with more selective sweeps of beneficial mutations. Previous research has also demonstrated that the spread of hypermutator alleles tends to be found in populations with a larger population size or weaker bottleneck effects (Raynes et al. 2014). As a result, even though the mutation rates of lines in two genetic backgrounds (WT and MMR-) evolved to be closer during the experiments with all three transfer sizes, the driving mechanisms are different between populations under the L transfer size and those under M or S transfer sizes (**Fig. 2**).

In theory, the evolved populations under larger transfer sizes experienced a relatively longer duration of stationary phase between transfers as the number of divisions needed to reach the stationary phase are smaller. Given that the physiological features of *E. coli* can be different in different growth phases (Pletnev et al. 2015), the evolutionary pressure under different transfer-sizes may also be different. Therefore, interpreting any results based on the comparison across transfer-size treatments needs more caution. In our case, however, such differences are likely limited, as we still found significant parallelism in the fixed enriched mutations across different transfer sizes (**Fig. S8**). Whether there is any adaptation specific to the different growth phases or different combinations of growth phases is a subject for future studies.

### The possible role of biofilm formation in adaptation

With respect to the specific genes and biological processes that appear to be targets for adaptation, our analysis of candidate genes suggests that biofilm formation is an important characteristic in adapting to the complex setting imposed by the experimental environment. In particular, the formation of type I fimbriae is critical for biofilm formation in *E. coli* (Pratt and Kolter 1998). Consistently, several genes enriched for fixed nonsynonymous mutations are related to formation of type I fimbriae (**Fig. 5B**), including *fimH* and *fimG*, which account for components the type I fimbriae (Waksman and Hultgren 2009; Le Trong et al. 2010), *fimB* and *fimE*, which regulate the expression of *fimAICDFGH* operon (Olsen et al. 1998), and *proQ* (RNA chaperone) and *lsrK* (autoinducer-2 kinase), which also facilitate biofilm formation (Li et al. 2007; Sheidy and Zielke 2013). We additionally observed an enrichment of mutations in the intergenic region of *fimE*/*fimA* (**Fig. 5C**) which contains a phase-variable promoter for regulating the expression of the *fimAICDFGH* operon (Abraham et al. 1985; Spears et al. 1986). Lastly, the list of genes enriched with structural mutations (**Fig. 5D**) also include *fimE*, which primarily turns off the expression of *fimAICDFGH* operon, and several genes related to *gatYZABCD* operon, including *gatZ*, *gatB*, and *gatA,* whose deletions can increase biofilm formation (Domka et al. 2007). Many of these candidate genes related to type I fimbriae contributed to the adaptation in an earlier experimental environment of a similar nature (Behringer et al. 2018). Moreover, structural mutations in *gatZ* and *gatA* genes have been found to contribute to the initial adaptation of *E. coli* in a mouse gut, another example of a complex environment (Barroso-Batista et al. 2014). Studying the genetic variants promoting the evolution of biofilm in complex environments may be of particular interests in the field of public health, as the evolution of biofilm has been considered to be related to the evolution of microbial social behaviors (Tarnita 2017), the evolution of pathogenicity (Kaper et al. 2004; Naves et al. 2008; Rossi et al. 2018), and the evolution of antibiotic resistance (Avalos Vizcarra et al. 2016; Sharma et al. 2016). Thus, understanding the evolution of biofilm formation may be key to increasing the efficiency of treatments in patients to combat the fast emergence of antibiotic resistance and pathogenicity.

### Candidate transcriptional regulators involved in adaptation

Because we have provided fresh media every day during the experimental evolution, a fast switch from stationary phase to exponential growth phase may bring benefits to the evolving populations (Monod 1949; Navarro Llorens et al. 2010). Interestingly, several genes enriched in fixed nonsynonymous mutations in our study are transcription regulators, including *arcA*, *cadC*, *cytR*, *rbsR*, *rseB*, and *sspA* (**Fig. 5B**). The genes enriched in structured variations also include transcription regulators, such as *arcB*, *cadC, rpoS,* and *nlpD* (**Fig. 5C**). The mutations on these genes may contribute to the transcriptomic reprogramming for the fast switch from stationary phase to exponential phase. For example, *rseB* is a negative regulator of the stationary phase effector, sigma factor E (Missiakas et al. 1997); and both *cytR* and *rbsR* are repressors to the carbon limitation effector, cAMP-CRP (Bell et al. 1986; Mauzy and Hermodson 1992; Kristensen et al. 1996). Therefore, gain-of-function mutations on these three genes can theoretically reduce the chance that cells stay in stationary phase. In addition, *arcA*, *arcB, cadC*, *rpoS,* and *sspA* are known as stress-responding activators (Iuchi et al. 1989; Lange and Hengge-Aronis 1991; Watson et al. 1992; Williams et al. 1994; Rolfe et al. 2011). Therefore, loss-of-function mutations on these genes are presumably beneficial in the experimental environment involving a frequent supply of fresh media and presumably imposing low stress. Further investigation will be needed to determine whether any of these mutations can bring such benefits to our experimentally evolved populations.

### Concluding Remarks

To sum up, our results reveal that high mutation rates in *E. coli* have only a very limited influence on the rate of adaptation. Our findings may provide useful insights for clinically relevant processes involving asexual populations, such as the evolution of improved growth rates in pathogens and the emergence of antibiotic resistance in natural or host environment. In particular, our experimental evolution results in complex media are likely to be more representative, as natural or host environments are usually highly heterogeneous. For example, a combination of the occasional emergence of hypermutators under weaker bottlenecks and the consistent evolution of antimutators under stronger bottlenecks may explain why a low to intermediate frequency of hypermutators is usually found in pathogen populations (Couce et al. 2016; Veschetti et al. 2020). Whether the elevated mutation rates affect adaptation rates and the pattern of genome evolution in these populations under natural or host environments should be subject to future research. For example, hypermutators may be critical for a founding population in a new environment, especially with epistasis in the fitness landscape, but such effect of hypermutation can diminish after a long time (Mehta et al. 2019).

## Materials and Methods

### Strains

The ancestral strains used in experimental evolution are descendants of PMF2 and PMF5, provided by the Foster Lab (Lee et al. 2012). PMF2 is a prototrophic derivative of *E. coli* K-12 str. MG1655, and its genetic background is called by WT in the paper. PMF5 is derived from PMF2 with *mutL* deletion, providing the MMR-genetic background. For both kinds of strains, a 3513 bp deletion to the *araBAD* operon is further introduced by lambda red recombineering as a neutral marker. Plates with TA agar (1% Arabinose, 1%Tryptone, 0.5% NaCl, 0.1% Yeast Extract, 0.005% TTC (Sigma T8877)) is used for examining the deletion of *araBAD* operon. The colonies with deletion (*ara*-) appear to be pink; otherwise, the colonies (*ara*+) appear to be purple.

### Experimental evolution

When we established the experimental populations, the ancestral strains were first cultivated overnight at 37 °C on LB agar plates, and then their single-isolated progenitor colonies were inoculated in a 16-× 100-mm glass tube with 10 mL of LB-Miller broth (BD Difco). The tubes were cultured at 175 rpm shaking at 37 °C. Every day, cultures are thoroughly-vortexed and transferred into a new tube with 10 mL of fresh LB broth. Three different transfer sizes are used: 1 mL(large), 1uL(medium), and 1nL(small), which correspond to different dilution factors: 10^-1^, 10^-4^, and 10^-7^. Initially, for each combination of genetic background and transfer sizes, we set up eight replicate tubes in three groups with different transfer sizes. For preventing cross-contamination, four replicates are *ara*- and four replicates are *ara*+. During the experimental evolution, experimental populations at day 90, 200, 300, 400, 500, 600, 700, 800, 900 were frozen in -80 °C freezers for analysis.

### Competition assay

When evaluating the fitness of an evolved population (*ara*-), we inoculated the corresponding frozen sample in a tube with 10 mL of fresh LB broth at 37 °C shaking at 175 rpm overnight. We also inoculated the corresponding ancestral population in a tube with 10 mL of fresh LB broth at 37 °C shaking at 175 rpm overnight. We then put 50 uL aliquot from evolved populations and 50 uL aliquot from evolved populations to into a new tube with 10 mL of LB broth at 37 °C shaking at 175 rpm for 24-hr competition. Immediately after inoculation (day 0) and after 24-hr competition (day 1), we used plate-counting to determine the colony-forming units (CFU) of both ancestral and evolved strains. Specifically, we serially diluted 100uL aliquot in 900 uL phosphate buffered saline. To distinguish the evolved population from the ancestors, we plated serially diluted aliquots on TA agar because *ara*-colonies will appear to be pink, while the ancestral (*ara*+) will appear to be purple. The TA plates were then incubated at 37 °C overnight. We then identified the plate with total colonies 30-300 and counted the number of colonies for the evolved population and the ancestors. Based on the colony numbers and the dilution factor during the serial dilution, we then calculated the CFU of ancestors before the competition (*A*0), the CFU of ancestors after the competition (*A*1), the CFU of the evolved population before the competition (*E*0), and the CFU of the evolved population after the competition (*E*1). The fitness of the evolved population (*w*) relative to the ancestor was then calculated by the following formula:

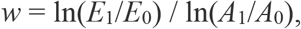

which is the ratio of two Malthusian parameters (Lenski et al. 1991).

At day 0, we also serially diluted and plated 100uL aliquot of evolved population as a control. For an evolved population, if both purple and pink colonies in the countable control plates (30-300 colonies per plate), its data will be discarded from the analysis because we are not sure about the source of the pink colonies in the experimental plates.

### Fluctuation test and mutation rate estimation

We quantified mutation rates of evolved populations (at day 900) and ancestors by fluctuation tests (Foster 2006). Briefly, fluctuation tests measure the rate of resistance to the antimicrobial rifampicin which is conferred by mutations to *rpoB*. For each combination of genetic background and transfer size, the four *ara*-populations were assayed. For each population, two biological replicates with different starting clones were assayed. For each of the WT or MMR-ancestor, we also run replicate experiments in different starting clones. For each clone, 40 replicate experiments were performed. Number of mutants as determined by CFU/mL were converted to an estimated mutation rate and a corresponding 95% confidence interval by the function “newton.LD” function in the R package “rSalvador”(Zheng 2017).

### DNA isolation and high-throughput sequencing

We conducted high resolution population tracking by collecting 1 mL of culture at day 90, 200, 300, 400, 500, 600, 700, 800, and 900 of experimental evolution. We used the DNeasy UltraClean Microbial Kit (Qiagen 12224; formerly MO BIO UltraClean Microbial DNA Kit) to extract DNA. For library preparation and sequencing, we submitted DNA to either the Hubbard Center for Genomic Analysis at the University of New Hampshire, the Center for Genomics and Bioinformatics at Indiana University, or the CLAS Genomics Facility at Arizona State University for library preparation and sequencing. Library preparation was done by the Nextera DNA Library Preparation Kit (Illumina, FC-121-1030) following an augmented protocol for optimization of reagent use (Baym et al. 2015) before being pooled and sequenced as paired-end reads on an Illumina HiSeq 2500 (UNH) or an Illumina NextSeq 500 (Indiana; ASU). The target depth is 100X.

### Sequencing analysis

We performed Sequencing analysis on the Mason and Carbonate high-performance computing clusters at Indiana University. The quality control of sequencing reads were performed by Cutadapt v.1.9.1 (Martin 2011), which removes residual adapters and trims low quality sequences. The qualified sequencing reads were then mapped to the *Escherichia coli* K-12 substr. MG1655 reference genome (NC_000913.3). All mutations and their derived allele frequencies (DAFs) were identified using Breseq v.0.30.2 with the predict-polymorphisms parameter setting (Deatherage and Barrick 2014). Furthermore, several criteria of further quality checks were applied to the samples which we will only include in the following analysis: (1) mean sequencing depths > 10; (2) any WT sample identified to contain the 1,830 bp deletion in *mutL* from the PMF5 progenitor strain was discarded; (3) regions lacking sequencing coverage (i.e. depth = 0) must be smaller than 5% of the genome; and (4) the sequencing result should reflect the correct genetic background in terms of *ara* markers, including a nonsynonymous SNP at position 66528, an intergenic SNP at position 70289, and a multiple base substitution mutation (SUB) at position 66533. For an *ara+* population, we required either of two SNPs showing DAF < 0.2. For an *ara-* population, we required either of two SNPs showing DAF > 0.8 or the SUB is detected.

In the end, 396 genomic profiles passed QC and were included in the following analysis (**Table S3**). In the case of M transfer size under WT background, only six out of eight replicates of evolved populations are left for the following analysis because the other two were potentially contaminated by MMR-strains. In the other five combinations of transfer size and genetic background, there are still eight replicates of evolved populations for the following analysis. For every evolved population subject to the following analysis, its sequencing profiles are available in at least seven different time points.

To make sure that we do not use the mutations that originated from the starting clone before experimental evolution in the analysis, we discarded any mutations with a DAF = 1.0 at one time point for at least 11 experimental populations with the same genetic background from the analysis. Furthermore, the highly repetitive sequences in *rsx* genes are known to cause errors in SNP calling (McCloskey et al. 2018), so they were also discarded from the analysis.

### Number of guaranteed generations

Given the observation that carrying capacity of experimental populations can be recovered within a transfer period (i.e. one day), the guaranteed generation numbers in 900 days for dilution factors = 10^-1^, 10^-4^, and 10^-7^ were respectively estimated as 900 times of log2(10), log2(10^4^), and log2(10^7^), which is equal to ∼ 3.0k, 12k, and 21k.

### Rate of genomic evolution

The level of genomic divergence for each experimental population at each time point is defined by summing all DAFs of detected mutations. We then calculated the mean genomic divergence across all eligible experimental populations in each combination of transfer size and genetic background. We further performed the linear regression by the function “lm” in R with formula “mean genomic divergence ∼ guaranteed generations + 0”, which enforces the y-intercept as 0. The slope of the regression is the estimated rate of genomic evolution. We also performed a nonlinear regression using the formula “mean genomic divergence ∼ guaranteed generations + square root of guaranteed generations + 0”, which was previously proposed to catch the trend of diminishing returns (Tenaillon et al. 2016).

### Identification of fixed mutations by hidden Markov chain

For each population, clade-aware hidden Markov model (caHMM) was performed using a modified version (Behringer et al. 2020) of previously released code (Good et al. 2017). For the populations in which caHMM can not finish, we instead performed well-mixed hidden Markov chain (wmHMM) using a modified version (Behringer et al. 2020) of previously released code (Good et al. 2017). The single clade in wmHMM is defined as the basal clade. Fixed mutations are then defined as mutations that are inferred to be fixed in basal, major, or minor clade in the results of either analysis.

### Estimation of selection coefficients

For each fixed mutation, we estimate its selection coefficients by its temporal data of DAFs. If a mutation is in the basal clade, no correction is needed. If caHMM infers a mutation belongs to the major or minor clade, its corrected allele frequency will be the DAF devided by the proportion of the population belonging to the major or minor clade (also inferred by caHMM). For each of two available consecutive time points *i* and *j*, if the corrected allele frequencies at both time points (*p*_i_ and *p*_j_) are smaller than 0.95 and larger than 0.05, the selection coefficient is calculated as

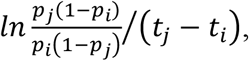

where *t*_i_ and *t*_j_ is the number of guaranteed generations at time point *i* and *j*, respectively. The negative values are discarded. The largest positive value across all pairs of time point are used as the final measurement.

### Calculation of neutrality index

As discussed in the main text, we defined (*F*_N_/*F*_S_)/(*U*_N_/*U*_S_) as the neutrality index of nonsynonymous SNPs, where *F*_N_ is the number of nonsynonymous SNPs fixed within a clade or fixed in an entire population, *F*_S_ is the number of synonymous SNPs fixed within a clade or fixed in an entire population, *U*_N_ is the number of nonsynonymous SNPs in the mutation-accumulation experiment, and *U*_S_ is the number of synonymous SNPs in the mutation accumulation-experiment. We also similarly defined (*F*_I_/*F*_S_)/(*U*_I_/*U*_S_) as neutrality index of intergenic SNPs, where *F*_I_ is the number of intergenic SNPs fixed within a clade or fixed in an entire population, and *U*_I_ is the number of intergenic SNPs in the mutation-accumulation experiment. The values of *U*_N_, *U*_S_, and *U*_I_ are from a previously published mutation-accumulation experiment of our ancestral lines (Lee et al. 2012). We calculated population-specific indexes and then acquired population-wise mean and SE. The populations with *F*_N_ = 0 and *F*_I_ = 0 are discarded in the calculation of neutrality index of nonsynonymous and intergenic SNPs, respectively.

### Calculation of *G*-scores

For each combination of genetic background and transfer size, we quantified the parallelism of the fixed nonsynonymous mutations using the sum of *G*-scores across genes (Tenaillon et al. 2016). A larger *G*-score for a gene suggests that the fixed nonsynonymous mutations are more overrepresented in that gene. Specifically, to calculate a genic *G*-score (*G*_i_), we first counted the observed number of fixed nonsynonymous mutations in gene *i* (*O*_i_) per a combination of genetic background and transfer size. We then calculated the expected number for gene *i* (*E*_i_) by *O*_tot_(*L*_i_/*L*_tot_), where *O*_tot_ *=* Σi *O*_i_, *L*_i_ is the number of nonsynonymous sites for gene *i,* and *L*_tot_ *=* Σi *L*_i_. In the end, *G*_i_ is defined by 2*O*_i_ln(*O*_i_ /*E*_i_) or defined as zero when *O*_i_ = 0 or when 2*O*_i_ln(*O*_i_ /*E*_i_) < 0.

It was noted that the null expectation of *G*-scores varies with total number of fixed nonsynonymous mutations (Behringer et al. 2020). Therefore, for each combination of genetic background and transfer size, we performed 20,000 simulations in each of which *O*_tot_ hits are randomly distributed among all *L*_tot_ sites across all genes in the reference genome. Then the significance of the sum of *G*-scores was evaluated by the *z* score defined by (the observed sum - mean of simulated sums) / (standard deviation of simulated sums).

### Calculation of mean Bray-Curtis similarity

For each combination of genetic background and transfer size, we also quantified the parallelism of the fixed nonsynonymous mutations using the mean Bray-Curtis similarity across all pairs of experimental populations for a TP/GB combination (Turner et al. 2018; Behringer et al. 2020). Specifically, for a pair of populations *j* and *k*, their Bray-Curtis similarity is defined by

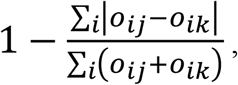

where *o*_ij_ and *o*_ik_ is the observed number of fixed nonsynonymous mutations in gene *i* for population *j* and *k*, respectively.

For each combination of genetic background and transfer size, we also performed 1,000 simulations to acquire the null distribution. In each simulation, we randomly sample the nonsynonymous sites up to the number of observed fixed nonsynonymous mutations for each population and calculated mean Bray-Curtis similarity as described above. After acquiring the null distribution, we evaluated the significance of the observed mean Bray-Curtis similarity by calculating the *z* score defined by (the observed value - mean of simulated values) / (standard deviation of simulated values).

### Overrepresentation of the genes affected by nonsynonymous mutations

To evaluate the significance of *G*-score for gene *i*, we directly compared the *G*_i_ to the distribution of 20,000 simulated *G*_i_, and the *P*-value was defined as the proportion of simulated *G*_i_ larger or equal to the observed *G*_i_. For multiple test correction, we multiplied each gene’s *P*-value by the number of genes with at least one hit by the set of fixed nonsynonymous mutations (Bonferroni correction). The genes are called significant only if the genes show Bonferroni corrected *P*-value < 0.05.

### Enrichment test of GO terms and KEGG pathways

Using the set of significant genes, we performed the enrichment test of gene ontology terms using the function “enrichGO” in R package “DOSE” (Yu et al. 2015) with *q*-value cut-off = 0.05 and the organismal database as org.EcK12.eg.db. We also performed the enrichment test of KEGG pathways using the function “enrichKEGG” in the same package but found no terms with *q*-value < 0.05.

### Overrepresentation of the intergenic regions affected by fixed mutations

The identification of the intergenic regions affected by mutations was also performed by the way similar to identify the genes affected by nonsynonymous fixed mutations in genic *G*-score approach (Tenaillon et al. 2016). Instead of focusing on genic regions, genome-wide intergenic regions are focused. For each combination of genetic background and transfer size, we first counted the observed number of intergenic mutations in intergenic region *i* (*O*_i_), and the expected number for intergenic region *i* (*E*_i_) was calculated by *O*_tot_(*L*_i_/*L*_tot_), where *O*_tot_ *=* Σi *O*_i_, *L*_i_ is the length for intergenic region *i,* and *L*_tot_ *=* Σi *L*_i_. The *G*-score for intergenic region *i* (*G*_i_) was then calculated by 2*O*_i_ln(*O*_i_ /*E*_i_), following the methods described in the above section. We also performed 20,000 simulations and determined the Bonferroni corrected *P*-value for each gene *i* following the methods described in the above section.

### Overrepresentation of the genes affected by structural fixed mutations

The identification of the genes affected by structural fixed mutations was performed by the way similar to identify the genes affected by nonsynonymous fixed mutations in genic *G*-score approach (Tenaillon et al. 2016). Structural mutations include indels and IS-element insertions. For each combination of genetic background and transfer size, we first counted the observed number of populations with any structural mutations in gene *i* (*O*_i_), and the expected number for gene *i* (*E*_i_) was calculated by *O*_tot_(*L*_i_/*L*_tot_), where *O*_tot_ *=* Σi *O*_i_, *L*_i_ is the gene length for gene *i,* and *L*_tot_ *=* Σi *L*_i_. The *G*-score for gene *i* (*G*_i_) was then calculated by 2*O*_i_ln(*O*_i_ /*E*_i_), following the methods described in the above section.

We also performed 20,000 simulations and determined the Bonferroni corrected *P*-value for each gene *i* following the methods described in the above section. As a result, we found all the genes with *O*_i_ ≥ 2 show Bonferroni corrected *P*-value < 0.05.

### Correlations between pairs of SNPs

For each evolved population, we focused on the nonsynonymous, synonymous, and intergenic SNPs in which at least two nonzero DAFs were found. For each pair of two such SNPs, we calculated the change of DAFs. Then we calculated Pearson’s correlation coefficients across only all the odd-numbered changes of DAFs to avoid non-independence (Lynch and Ho 2020). That is to say, if a population has sequencing profiles available for analysis at every sampling time points (days 90, 200, 300, 400, 500, 600, 700, 800, 900), we will calculate Pearson’s correlation coefficients using the five changes of DAFs: the one between day 0 and day 90, between day 200 and day 300, between day 400 and day 500, between day 600 and day 700, and between day 800 and day 900. For another example of population, if its sequencing profile at day 90 is discard from analysis due to low quality, the Pearson’s correlation coefficients using only the four changes of DAFs: the one between day 0 and day 200, between day 300 and day 400, between day 500 and day 600, between day 700 and day 800. Note that at least four changes of DAFs are used for each evolved population because no populations have more than two missing profiles. We then get the distribution of Pearson’s correlation coefficients for each evolved population.

To establish the baseline for comparison in each combination of transfer size and genetic background, we also generated the set of Pearson’s correlation coefficients using two random mutations from two different random populations with all sequencing profiles available (i.e. they are unlinked for sure). We followed the same procedure above to calculate Pearson’s correlation coefficients for each pair of unlinked mutations. When simulating a distribution of Pearson’s correlation coefficients, we used 100 pair of unlinked mutations. We then repetitively performed 100 rounds of simulation to get the mean and SE for the distribution.

### Data Availability

Sequencing data generated during this study are available at NCBI’s Sequencing Read Archive: BioProject PRJNA532905 and PRJNA722381.

### Code Availability

Codes generated to analyze sequencing data are available at https://github.com/LynchLab/ECEE_Hypermutator

## Acknowledgements

We thank D. Allan Drummond, Jay T. Lennon, and James B. McKinlay for their helpful comments. High Performance Computing Resources were provided and maintained by the National Center for Genome Analysis Support at Indiana University. This work was supported by Army Research Office Grants ARO65308-LS-MU and W911NF-14-1-0411 (M. L.) and National Institutes of Health Grant F32GM123703 (M. G. B.) and R35GM122566 (M. L.).

## Author Contributions

Conceptualization, W-C.H., M.G.B., and M.L.; Methodology, W-C.H., M.G.B., and S.F.M.; Investigation, W-C.H., M.G.B., S.F.M., J.G., A.N., M.A., and G.F.B.; Formal Analysis, W-C.H., and M.G.B; Writing – Original Draft, W-C.H, and M.G.B.; Writing – Review & Editing, W-C.H., M.G.B., W-C.H., and M.L.; Visualization, W-C.H. and M.G.B.; Funding Acquisition, M.G.B. and M.L.

